# Decoupling diffusion, turnover, and advection in long-term FRAP

**DOI:** 10.1101/2025.02.04.636435

**Authors:** Outa Nakashima, Takumi Saito, Shinji Deguchi

**Author notes:** Corresponding author Address: 1-3 Machikane-yama, Toyonaka, Osaka 560-8531, Japan.

## Abstract

Intracellular molecular turnover is a dynamic process governed by diffusion, biochemical reactions, and intracellular transport dynamics. While fluorescence recovery after photobleaching (FRAP) has been widely used to quantify fluorescence recovery mechanisms, conventional models primarily focus on diffusion and reaction kinetics, often overlooking the influence of intracellular advection. However, in cytoskeletal structures such as stress fibers, myosin-driven actin retrograde flow generates a significant advective component, which complicates the interpretation of FRAP data, particularly in long-term observations where advective effects become sufficiently pronounced. Here, we develop an analytical framework that extends FRAP modeling to incorporate the coupled effects of diffusion, turnover, and intracellular advection within the photobleached region of interest. By deriving exact solutions to a reaction-diffusion-advection system, we identify three key dimensionless parameters that govern fluorescence recovery dynamics: the turnover-to-diffusion ratio, the monomer-to-filament ratio, and the advection magnitude. Our results demonstrate that, even in the absence of biochemical reactions, fluorescence recovery in a fixed region can occur due to advection, leading to potential misinterpretations of molecular exchange rates. The model provides a theoretical foundation for distinguishing these effects and offers a practical tool for long-term FRAP analysis, where the interplay of diffusion, turnover, and advection becomes increasingly relevant over extended timescales. By systematically characterizing the interplay between molecular diffusion, reaction kinetics, and intracellular transport, our framework provides deeper insight into protein turnover in complex biological environments.

## Introduction

Intracellular structures undergo continuous molecular turnover, enabling cells to adapt to applied stress (Kaunas & Deguchi, 2011; Ueda, Matsunaga, & Deguchi, 2024). The mechanisms driving turnover vary depending on the timescale (Saito, Matsunaga, & Deguchi, 2022b). Short-term turnover, primarily governed by diffusion and chemical association/dissociation, occurs typically within seconds to minutes. In contrast, long-term turnover involves additional factors such as advection. Often overlooked, advection can significantly influence turnover measurements. Specifically, bulk flow becomes relevant when proteins closely bound to actin cytoskeletal structures are translocated by myosin-driven and actin polymerization-induced retrograde flow (Gardel et al., 2008; Yamashiro et al., 2023; Yolland et al., 2019). This contribution of advection complicates the interpretation of long-term turnover measurements.

Fluorescence recovery after photobleaching (FRAP) is a widely used technique for evaluating protein turnover. While single-exponential fitting is commonly applied to analyze the recovery of fluorescent proteins, more complex scenarios require diffusion-reaction equations to distinguish between the contributions of diffusion and chemical association/dissociation. Traditional FRAP models assume spatially uniform and immobile chemical equilibrium distributions, limiting their applicability to short-term observations (McGrath, Tardy, Dewey, Meister, & Hartwig, 1998; Tardy, McGrath, Hartwig, & Dewey, 1995). For long-term FRAP analyses, where single-exponential fitting can become inadequate, double-exponential models are often employed (Campbell & Knight, 2007; Sakurai-Yageta, Maruyama, Suzuki, Ichikawa, & Murakami, 2015). However, the interpretation of the two independent time constants in double-exponential models remains unclear particularly concerning the actual physicochemical factors such as diffusion, chemical association/dissociation, and advection.

To address these complexities, we previously developed a numerical tool capable of automatically tracking the region of interest (ROI) during FRAP measurements (Saito, Matsunaga, & Deguchi, 2022a). This tool enables the separate analysis of diffusion, chemical equilibrium constants, and intracellular flow velocity. Despite its effectiveness, automatic ROI tracking is not always feasible for researchers who conventionally fix the ROI position. Fixed ROI measurements, while widely used in many studies (Hotulainen & Lappalainen, 2006; Katsuta et al., 2023), are susceptible to errors introduced by advection, leading to changes in fluorescence signals even when diffusion and association/dissociation are minimal. Consequently, developing a model that accurately interprets conventional FRAP data with a fixed ROI is crucial, especially for long-term observations of actin-based structures. Here, we present a FRAP analytical model developed for fixed ROI analysis, accounting for myosin-driven actin retrograde flow, namely intracellular advection.

## 2. Analytical model

### 2.1. Governing equations

The system of interest within a cell is represented as a rectangular structure with constant thickness and uniform properties (Fig. 1). The spatial coordinate *x* denotes the position along the system of length *L*, with the photobleached band of width *w*. Considering a sufficiently elongated cell, where intracellular flow is predominantly in the longitudinal direction, the problem can thus be effectively reduced to one dimension. Monomeric actin is modeled as belonging either to the filamentous meshwork or to the diffusive pool. The model assumes no net change in the total amounts of fluorescent and non-fluorescent actin molecules, nor in the relative proportions of filamentous meshwork and diffusive pool, with polymerization and depolymerization balanced throughout the system over time *t*. Under these conditions, the time-series changes in the concentrations of fluorescent monomers, categorized into diffusive monomers (*c*_*m*_) and those incorporated into the filamentous meshwork (*c*_*f*_), are governed by diffusion with a diffusion coefficient 𝒟, turnover at a rate *k*, and advection with a velocity of *u*:

**Fig. 1.**
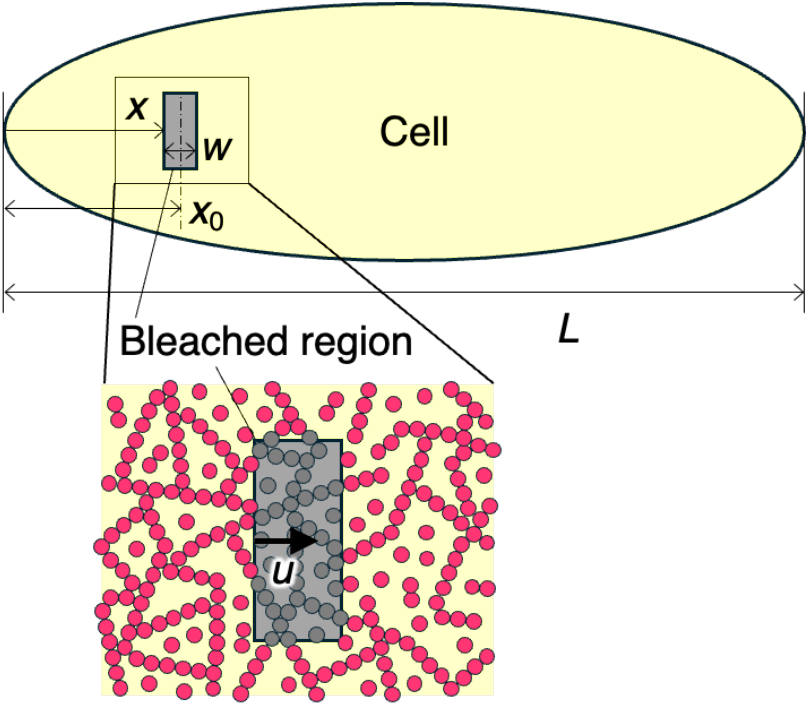
Schematic representation of the cellular system, consisting of monomeric actin (red particles) and a filamentous meshwork (polymerized actin filaments), with the fixed bleached region advected by a flow of velocity *u*.

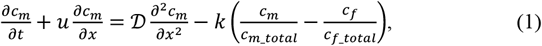

and

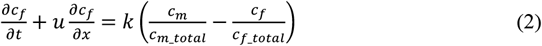

where *c*_*m*_*total*_ and *c*_*f*_*total*_ are the total concentrations of fluorescent monomers within the diffusive pool and filamentous meshwork, respectively. The rationale behind the reaction terms of the exchange process in the above reaction-diffusion-advection equations is as follows. The probability of monomers departing from the filamentous meshwork is expressed as *c*_*f*_/*c*_*f*_*total*_, while that from the diffusive pool is given by *c*_*m*_/*c*_*m*_*total*_, and the two groups exchange at a constant rate. The initial conditions are given by

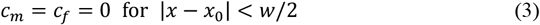

and

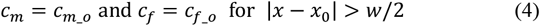

where *x*_*0*_ is the center of the bleached region, and *c*_*m*_*ext*_ and *c*_*f*_*ext*_ are the concentrations of fluorescent actin in the diffusive pool and filamentous meshwork outside of the bleached region, respectively.

### 2.2. Nondimensionalization

The above governing equations and initial conditions are nondimensionalized using the following dimensionless numbers marked with an asterisk

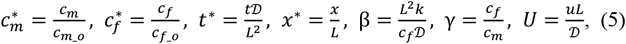

and the resulting equations are

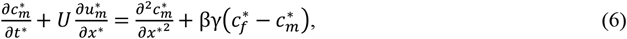

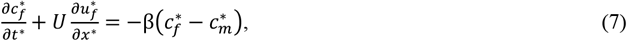

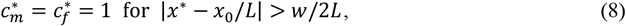

and

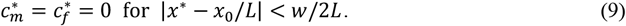

### 2.3. Analytical solutions

Assuming periodic boundary conditions, the variables are expressed using Fourier coefficients for the wavenumber *ξ*,

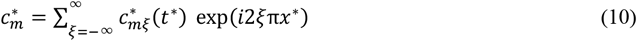

and

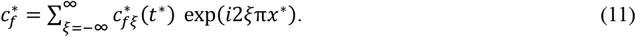

Substitution of Eqs. (10) and (11) into Eq. (6) and (7) yields

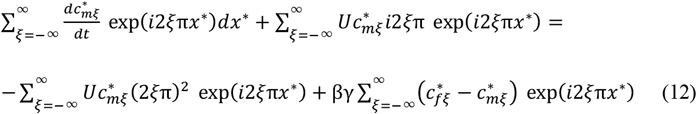

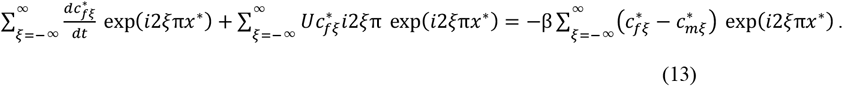

Multiplying these equations by exp(− *i*2*l*π*x*^∗^) and integrating from 0 to 1 extract the component for the wavenumber *l* as follows:

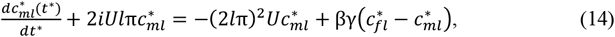

and

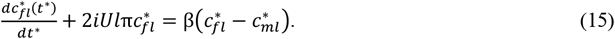

The resulting equations are transformed using the Laplace transform expressed in the *s*-domain and are organized into a matrix form

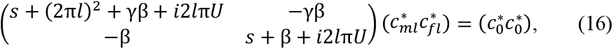

and hence the solutions are

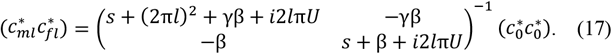

Applying the inverse Laplace transform yields

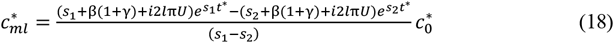

and

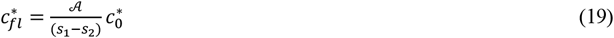

where

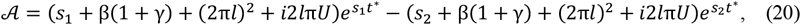

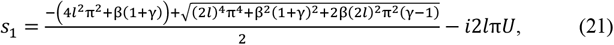

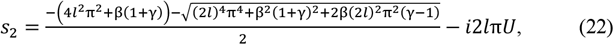

and

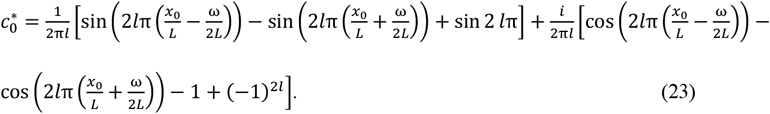

Equations (18) and (19) are rewritten with alternative variables to simplify their notations as follows

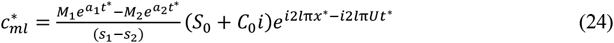

and

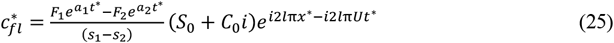

where *S*_0_ and *C*_0_ are even and odd functions of the wavenumber *l*, respectively,

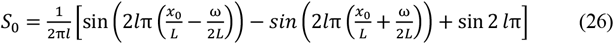

and

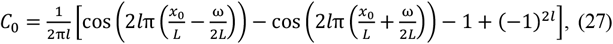

and

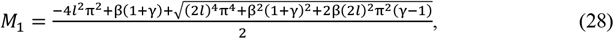

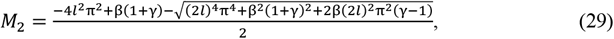

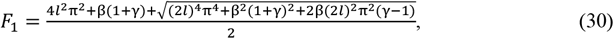

and

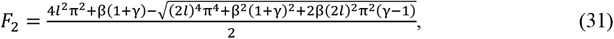

and *a*_1_ and *a*_2_ are the real parts of *s*_1_ and *s*_2_, respectively,

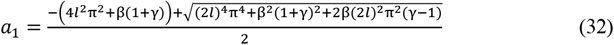

and

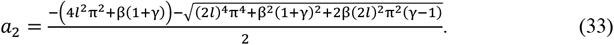

Considering the properties of odd and even functions, the final solutions are obtained as follows:

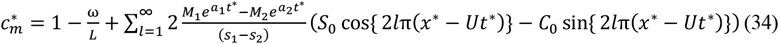

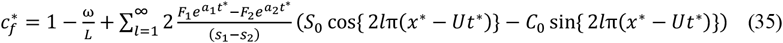

These analytical solutions, describing the time evolution of fluorescence recovery for actin monomers in the diffusive pool and filamentous meshwork, respectively, capture the effects of diffusion, turnover, and advection as a function of the three parameters: *β* (turnover-to-diffusion ratio), *γ* (monomer-to-filament ratio), and *U* (advection magnitude). Note that these equations satisfy the periodic boundary conditions:

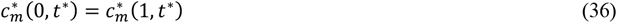

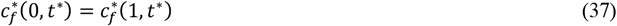

### 2.4. Net FRAP dynamics

The weighted average *F*, obtained from the proportions of molecules belonging to each group, is normalized by the maximum fluorescence *F*_*0*_ to provide the overall fluorescence recovery:

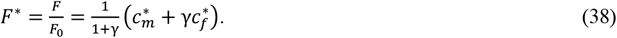

By integrating this expression over the bleached region and determining the mean intensity, the time evolution of fluorescence recovery within the fixed bleached area is obtained

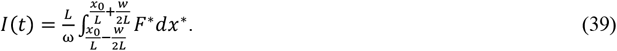

The integrated analytical solutions for the diffusive monomer group and the filamentous meshwork group are shown by

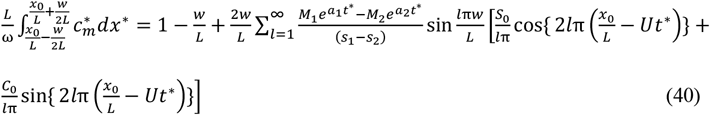

and

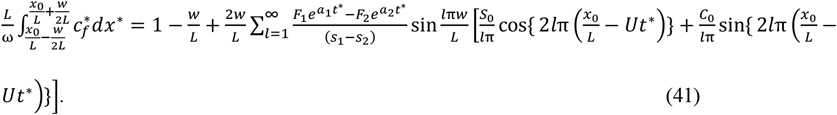

## 3. Results and discussion

The present study aimed to address challenges in extracting long-term intracellular protein turnover from conventional FRAP data. While short-term FRAP behavior within cells is primarily governed by diffusion and turnover, long-term turnover involves an additional influential factor, advection, driven by myosin-actin retrograde flow. Most existing FRAP models assume immobile equilibrium distributions (Campbell & Knight, 2007; Ciocanel, Kreiling, Gagnon, Mowry, & Sandstede, 2017; Hallen, Liang, & Endow, 2008; Saito, Kikuchi, & Ishikawa, 2024; Saito, Matsunaga, Matsui, Noi, & Deguchi, 2021; Sakurai-Yageta et al., 2015; Sprague et al., 2006), limiting their applicability to short-term analyses. Specifically, under sufficient advection, apparent fluorescence recovery in a fixed ROI can occur even in the absence of diffusion or turnover (Saito, Matsunaga, & Deguchi, 2022b). This phenomenon arises when non-bleached fluorescent structures from adjacent regions flow into the ROI, resulting in an increase in fluorescence intensity. Although unrelated to diffusion or turnover, it can lead to the misleading interpretation that turnover has occurred. To address this issue, we previously developed a dynamic ROI-tracking model capable of isolating each contributing factor (Saito, Matsunaga, & Deguchi, 2022a). However, many FRAP users may prefer measurements with a fixed ROI due to its simplicity and widespread usage.

Here, we developed a fixed ROI-based model, providing a framework for simultaneously detecting the individual effects of diffusion, turnover, and advection from a single FRAP measurement. The recovery behavior is analytically derived, enabling the interpretation of complex multiphysicochemical phenomena through three dimensionless parameters, β (turnover-to-diffusion ratio), γ (monomer-to-filament ratio), and U (advection magnitude), which quantify the relative contributions of each process. While intracellular dynamics has been theoretically interpreted through analytical models (Kang, Day, Di Benedetto, & Kenworthy, 2010; Kubitscheck, Wedekind, & Peters, 1994; Lele & Ingber, 2006; Lopez, Dupou, Altibelli, Trotard, & Tocanne, 1988; Sbalzarini, Mezzacasa, Helenius, & Koumoutsakos, 2005; Seiffert & Oppermann, 2005; Tardy et al., 1995; Wedekind, Kubitscheck, Heinrich, & Peters, 1996), our approach overcomes the challenges associated with fixed ROI-based FRAP experiments, offering a practical tool for analyzing intracellular turnover over extended timescales.

Stress fibers are dynamic structures that continuously undergo assembly and disassembly, playing critical roles in adaptive cellular remodeling (Kassianidou & Kumar, 2015; Ueda, Matsunaga, & Deguchi, 2022; Ueda et al., 2024), the regulation of pro-inflammatory responses (Huang et al., 2021; J. D. Humphrey, 2007; Jay D. Humphrey, Dufresne, & Schwartz, 2014; Kaunas & Deguchi, 2011; Saito, Huang, et al., 2021), and the progression of senescence (Chantachotikul, Liu, Furukawa, & Deguchi, 2025; Liu, Matsui, Kang, & Deguchi, 2022). These processes are driven by the coordinated activity of signaling and structural molecules. Understanding the mechanisms behind these processes requires precise analysis of molecular turnover, with a clear distinction between the contributions of diffusion, turnover, and advection. However, as these phenomena often occur simultaneously, the lack of FRAP models that adequately account for the influence of advection has made their differentiation challenging. For example, stress fibers, primarily composed of actin, myosin, and various associated proteins, are subject to retrograde flow. This advection influences apparent turnover in fixed ROI-based FRAP analysis, complicating efforts to accurately resolve the true dynamics of these structures.

Further studies should integrate detailed analyses with experimental approaches to refine our understanding of intracellular turnover mechanisms.

